# A new thalamo-cortical-amygdala circuit is involved in processing a natural auditory alarm cue

**DOI:** 10.1101/716910

**Authors:** Ana G Pereira, Matheus Farias, Marta A. Moita

## Abstract

Animals use auditory cues generated by defensive responses of others to detect impending danger. Here we identify a neural circuit in rats involved in the detection of one such auditory cue, the cessation of movement-evoked sound resulting from freezing. This circuit comprises the dorsal sub-nucleus of the medial geniculate body (MGD) and downstream areas, the ventral area of the auditory cortex (VA) and the lateral amygdala (LA). This study suggests a role for the auditory offset pathway in processing a natural sound cue of threat.

## INTRODUCTION

The use of auditory alarm cues from conspecifics is widespread across the animal kingdom^1–4^. Most research has focused on actively emitted signals, such as alarm calls and foot stamping^1,3^. However, auditory cues generated by movement patterns of prey can play a crucial role in predator-prey interactions. While it is well known that predators use motion cues produced by moving prey for their detection^5–8^, less is known about the ability of prey to use these motion cues to detect impending danger. Nevertheless, it has been established that crested pigeons use distinct wing whistles produced by conspecific escape flights^2^ and rats use silence resulting from freezing, as alarm cues^4^. Recently, it has also been suggested that seismic waves produced by fast running in elephants promote vigilance in conspecifics^9^.

The neuronal pathways underlying defensive responses triggered by sounds have been extensively studied using classical conditioning to pure tones, sweeps or white noise^10^. However, little is known about the mechanisms by which natural sounds, that most likely shaped the auditory system through evolution, trigger defensive responses^11,12^. Previously, we found that rats use freezing by others as a cue of danger. Importantly, the cessation of the movement-evoked sound resulting from the onset of freezing was the alarm cue. In playback experiments, this cue was sufficient to trigger freezing in rats^4^. Hence, in the present study, we probed the neuronal circuits involved in the processing of natural sound cues of threat by testing the response of rats to the transition from movement-evoked sound to silence.

## RESULTS

First, we exposed individual rats to unsignaled footshocks, as prior studies found that rats previously exposed to shock but not naïve ones respond to the distress of others^4,13^. The following day we tested their freezing response to the cessation of movement-evoked sound (Silence test) (Fig. 1a). During the test, rats were placed in a box and allowed to explore for three minutes during which a speaker played the recorded sound of another rat moving^4^ (see Methods). After this baseline period the sound ceased for one minute (silence period). We started by looking at the role of the lateral nucleus of the amygdala (LA), broadly implicated in freezing driven by learned aversive sounds^10^. To this end we optogenetically inactivated the LA using ArchT, a genetically encoded proton pump that hyperpolarizes neurons upon illumination with green light^14^. Importantly, the inactivation started 10 seconds before the onset of silence to encompass the transition from the movement-evoked sound to silence (Fig. 1c and Methods). In this experiment we used three groups: ArchT+light where neuronal activity is manipulated; and 2 control groups, Control light and Control ArchT, where neurons were undisturbed (see Methods). Since there was no significant difference in freezing between the two control groups (Supplementary Fig. 1b) we combined them into a single Control group. During the Silence test we found virtually no freezing during the baseline period in both ArchT+light and Control animals. Silence onset led to a robust increase in freezing in the Control group while in the ArchT+light there were minimal changes in rats freezing behaviour (Fig. 1c and 1d and Supplementary Figure 1c, 1d and 1e). This result shows that activity in the LA is necessary for the display of freezing triggered by the transition from movement-evoked sound to silence.

**Fig. 1.**
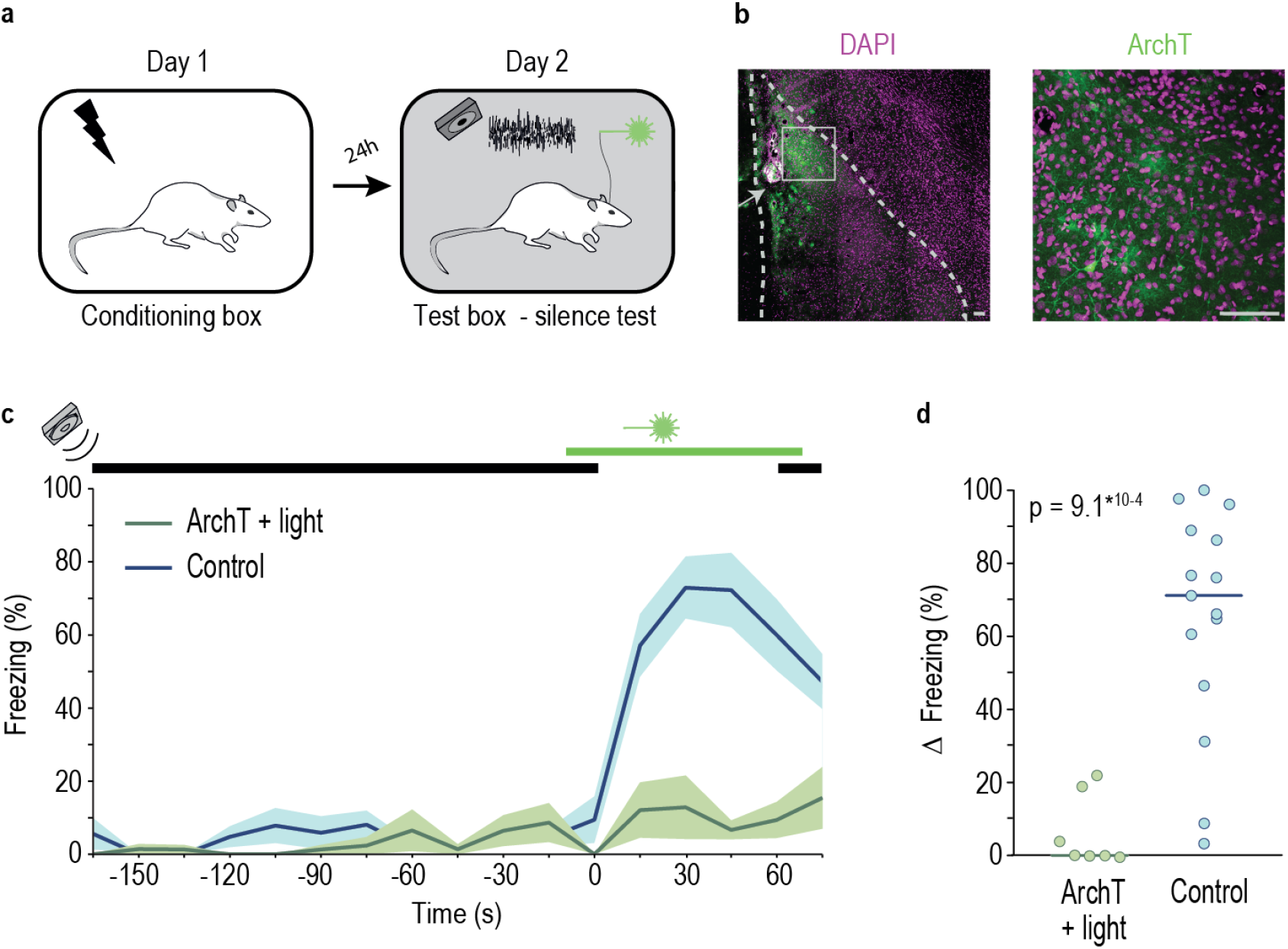
Optogenetic inactivation of the LA abolishes freezing triggered by the cessation of the movement-evoked sound. **a**, Schematics of the behavioral paradigm. **b**, Left - image of a coronal brain section from a representative rat, in which neurons in LA expressed ArchT; Right – image depicting a higher magnification of the area highlighted by ☐ on the left image. ↑ indicates tip of the injector. Scale bar 100μm. **c**, Movement-evoked sound (indicated by black bar) was played to rats from the ArchT + light (green, n=7) and Control (blue, n=15) groups with an interval of one minute of silence (between 0 and 60 seconds). Green bar indicates the period during which 556nm light was on. Line graph shows average % of freezing throughout the test session (shaded area shows dispersion of the data given by +/− sem). **d**, Individual dots represent the change in the percentage of time each animal spends freezing during the minute immediately preceding the cessation of the movement-evoked sound and the minute of silence. Horizontal bar represents the median value of the group (ArchT + light = 0.00%, Control = 71.20%, Wilcoxon Rank Sum test).

The LA receives auditory information through a direct pathway originating in several sub-nuclei of the medial geniculate body (MGB) of the auditory thalamus^15^. These same MGB sub-nuclei are also the origin of a indirect pathway that involves connections through the auditory cortex as well as other cortical regions^10,15,16^. To study the auditory pathways that convey information about the cessation of movement-evoked sound to the LA, we started by asking which of MGB’s sub-nuclei projecting to LA both directly and indirectly respond to the cessation of movement-evoked sound (Fig. 2a). To this end, we quantified the expression of the neural activity marker *c-fos* in 2 groups of animals - one exposed to continuous playback of sound and another exposed to the same sound but with 2 periods of silence (see Methods). As expected, we found that rats in the silence group increased freezing during the two one-minute silence period relative to the preceding minute (median change in freezing = 40.27%). No increase was observed when comparing the same time periods for rats in the continuous sound group (median change in freezing = −1.53%; Wilcoxon Rank Sum test comparing median change in freezing of silence and continuous sound groups, p = 0.0303, ranksum = 53.5). The quantitative analysis of *c-fos* expression revealed that exposure to silent periods significantly increased the number of *c-fos* positive cells in the dorsal division of the MGB (MGD) in comparison to continuous sound exposure (Fig. 2b). In addition, we found that the number of *c-fos* positive cells increased from anterior to posterior regions of the MGD (Fig. 2d). Interestingly, prior electrophysiology studies in anaesthetised rodents showed a higher prevalence of sound offset responses in the MGD relative to other sub-nuclei of the auditory thalamus^17^ and higher number of offset cells in the caudal part of MGD^18^. Together, these findings suggest that the cessation of the movement-evoked sound triggers sound offset responses in the MGD, which can then drive LA neurons either directly or indirectly.

**Fig. 2.**
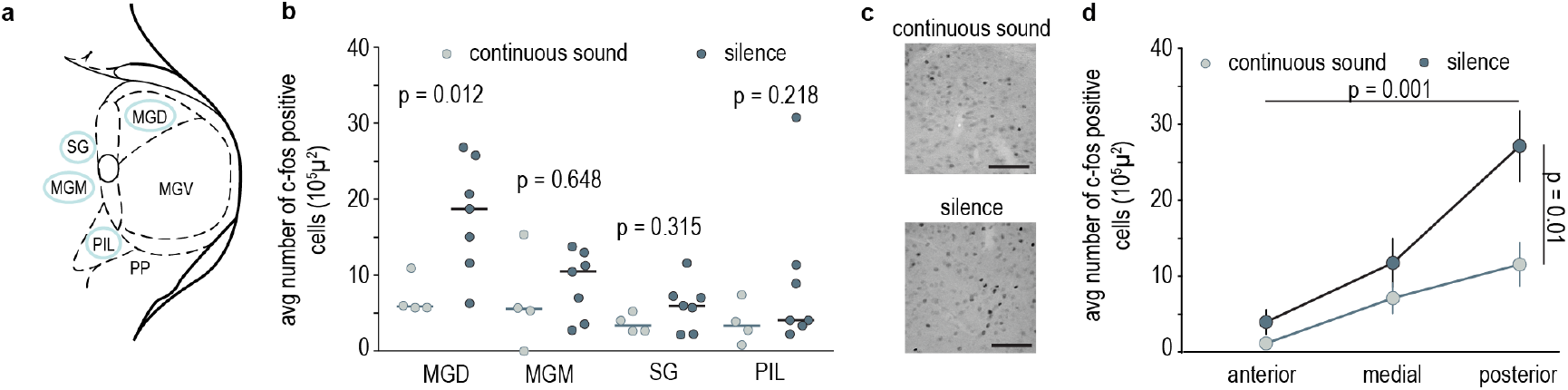
Differential activity in the dorsal division of the MGB triggered by the cessation of movement-evoked sound. **a**, Schematics of the MGB and its subnuclei. Regions circled in blue are part of both the mono and polysynaptic pathway to the LA. **b**, Average number of *c-fos* positive cells in the dorsal (MGD) and medial (MGM) divisions of the MGB, the suprageniculate nucleus (Sg) and the posterior intralaminar thalamic nucleus (PIL) of individual rats from the continuous sound (light grey n=4) and silence (dark grey n=7) group. Horizontal bar represents the median value and Wilcoxon Rank Sum test was used for comparisons (MGD continuous sound = 5.87, silence = 18.7, rank sum = 11; MGM continuous sound = 5.54, silence = 10.50, rank sum = 21; Sg silence = 5.95, continuous sound = 3.36, rank sum = 18; PIL silence = 4.07, continuous sound = 3.33, rank sum = 17). **c**, Representative pictures of c*-fos* labelled cells in MGD. **d**, Line graph showing average number of *c-fos* labeled cells in the MGD along the Anterior-Posterior (AP) axis for continuous sound and silence groups. Vertical bars represent +/− sem (*n*-way ANOVA main effect repeated measures AP axis p = 0.001, F = 12.69; silence vs continuous sound effect p = 0.01, F = 7.71).

Next, we tested whether MGD activity is required for silence-triggered freezing by optogenetically inactivating this area (Fig. 3a). We found that freezing was greatly reduced upon MGD inactivation relative to controls (Fig. 3d and 3e and Supplementary Figure 2c, 3d and 3e). Several studies addressed the anatomical and physiological properties of the MGD^15,17–19^. However, to our knowledge, the contribution of this sub-nucleus to behaviour remained untested. Therefore, we asked whether the MGD is required for freezing in response to a conditioned tone, as are other MGB sub-nuclei that project to LA^20^. To this end, we conditioned the same rats used in the previous experiment with three tone-shock pairings. The following day we tested their response to the conditioned tone while optogenetically inhibiting the MGD (Fig. 3a). Inactivation of the MGD did not affect the expression of freezing triggered by the conditioned pure tone (Fig. 3e). These results show that activity in the MGD is necessary for the display of freezing triggered by the transition from movement-evoked sound to silence but not that triggered by a conditioned pure tone. In line with physiological data, this indicates that the MGD may be preferentially recruited to process the offset of sound^17,18^.

**Fig. 3.**
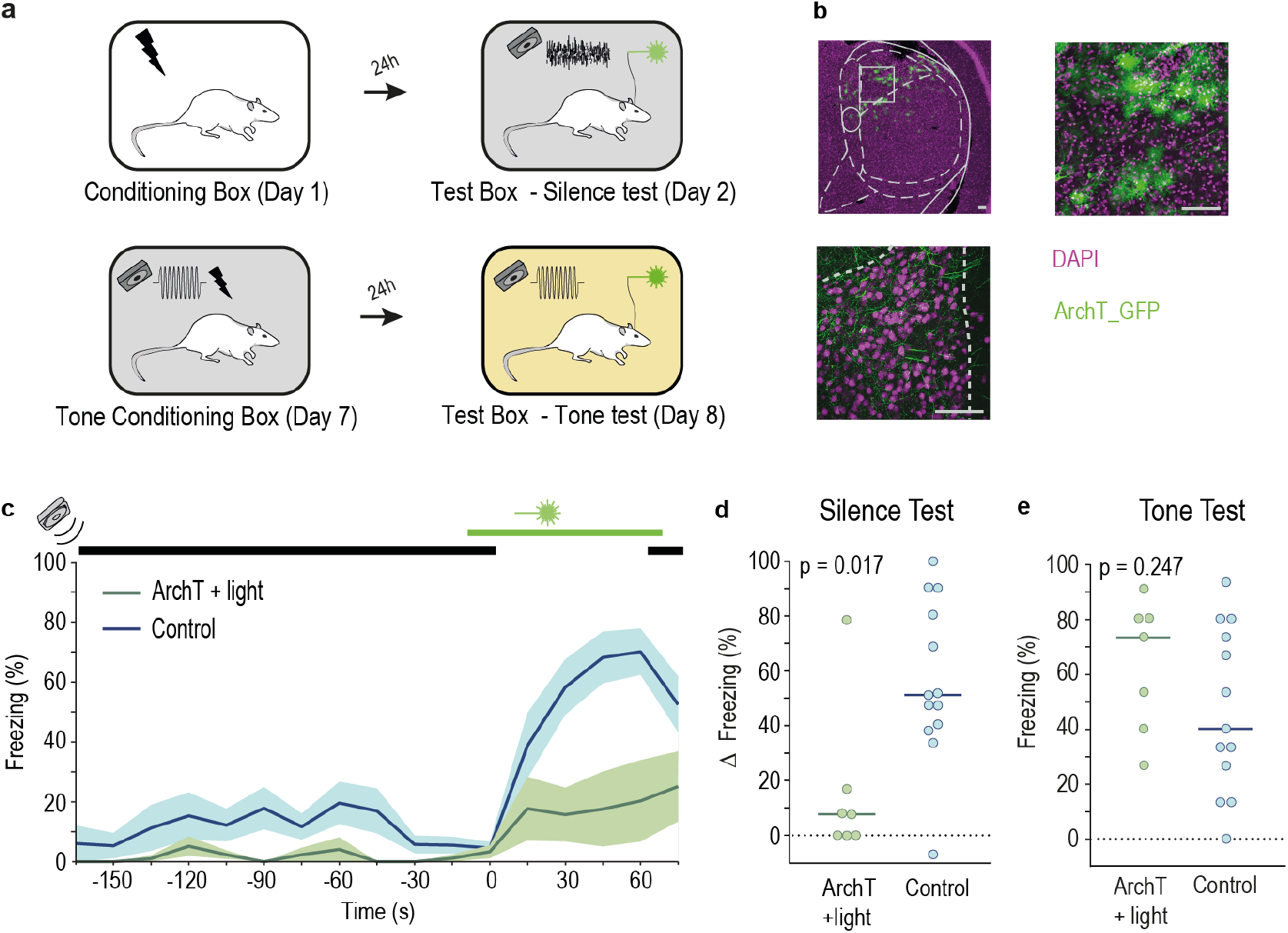
Activity in the dorsal division of the MGB is necessary for freezing triggered by the cessation of movement-evoked sound but not by a conditioned tone cue. **a**, Schematics of the behavioral paradigm used for testing the role of the MGD in freezing triggered by the cessation of movement-evoked sound and 5KhZ pure tone. **b**, Top Left - coronal brain section from a representative rat, in which MGD neurons express ArchT; Top Right – image with a higher magnification of the area highlighted by ☐ on the left image. Bottom - axon terminals in the tip of LA originating from ArchT expressing cells in MGD. **c and d**, Same as Fig.1c and 1d (respectively) for rats expressing ArchT and exposed to light illumination in the MGD (n = 7) and respective controls (n = 13). Horizontal bar represents the median value (ArchT + light = 7.73%, Control = 51.20%, Wilcoxon Rank Sum test, rank sum = 43). **e**, Individual value plot displaying the change in the percentage of time animals spent freezing during the 15sec immediately preceding the presentation of the 5KHz conditioned tone and the 15sec during which the tone was played. Horizontal bar represents the median value (ArchT + light = 73.33%; Control = 40.00 %; Wilcoxon Rank Sum test P = 0.247, ranksum = 88.5). Scale bar 100 μm.

The MGD sends direct projections to the LA^15^ but also projects to discrete areas in the cortex that through polysynaptic connections can convey information to the LA. To test if auditory processing areas downstream of the MGD are necessary for the response to the cessation of the movement-evoked sound, we focused on the two regions in the temporal cortex to which MGD axons project to - posterodorsal (PD) and ventral area (VA)^19,21^. We inactivated PD or VA using a pharmacological approach (Supplementary Fig. 3a and 3b). Briefly, rats with bilateral cannulas targeting PD or VA received a microinjection of the GABA agonist muscimol, just prior to the silence test (see Methods). At the end of the experiment animals received an injection of the retrograde tracer cholera toxin B (CTB), which allowed confirmation that the inactivated cortical region received input from MGD (Fig. 4c and 4f). Inactivation of PD with muscimol did not significantly decrease freezing triggered by silence in comparison with control rats (Fig. 4a and b). Inactivating VA, however, strongly impaired freezing during the silence period (Fig. 4d and 4e). These results show that activity in VA, but not PD, is necessary for freezing triggered by the cessation of movement-evoked sound.

**Fig. 4.**
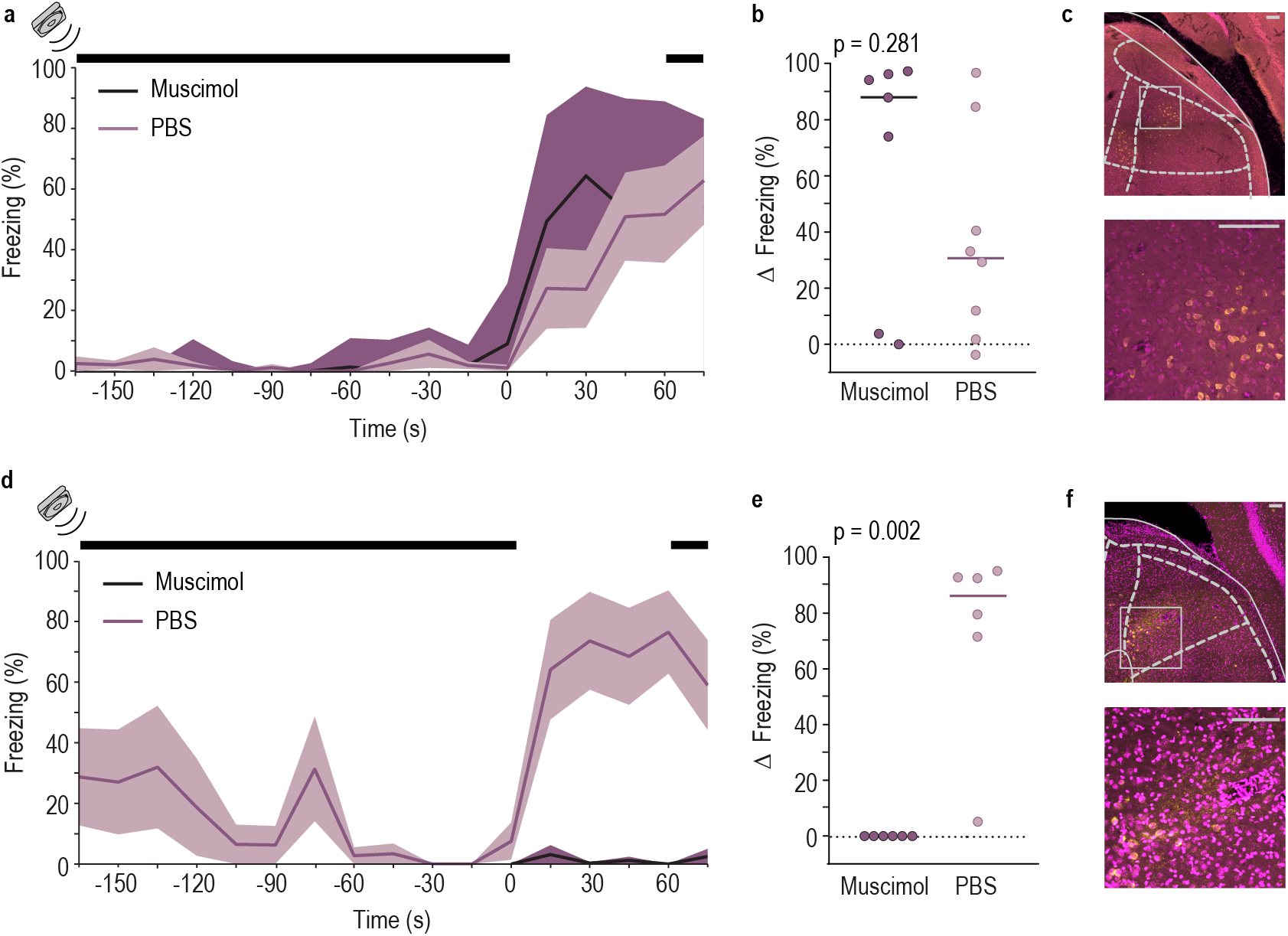
The ventral area (VA) but not the posterodorsal (PD) region of the auditory cortex is necessary for freezing triggered by silence. **a**, and **b**, Same as Fig. 1c, and d, (respectively) but for rats with bilateral injection of Muscimol or PBS in the area PD. Horizontal bar represents the median value (Muscimol = 87.87%, PBS = 30.73%, Wilcoxon Rank Sum test, ranksum = 54). **c**, Top - coronal brain section from a representative rat in which MGD neurons are labelled with Cholera Toxin subunit B injected in the PD, Bottom – image with higher magnification of the area highlighted by ☐ on the top image. **d**, and **e**, Same as **a**, and **b**, (respectively) but for rats with bilateral injections in VA. Horizontal bar represents the median value (Muscimol = 0.00%, PBS = 86.07%, Wilcoxon Rank Sum test, ranksum = 57). **f**, Same as **c**, but for Cholera Toxin subunit B injection in the VA.

## DISCUSSION

Together, our results suggest that the MGD, VA and LA form a network important for processing the transition from movement-generated sound to silence. Freezing triggered by the cessation of movement sound is likely to rely on cells with sound offset responses, although further experiments are necessary to confirm this hypothesis. Reports demonstrating the presence of sound-offset neurons in the MGD and their output regions in the auditory cortex^17,18,22–24^ support our proposed network. Indeed, a growing body of work has identified neurons with sound-offset responses throughout the auditory system. These reports suggest the existence of an “offset pathway” thought to be of major importance in processing speech, sound localization and movement detection^25^. Importantly, in most of these studies the offset responses are transient, lasting a fraction of a second, while in our study freezing lasts for several seconds. One possibility is that cells with offset responses act as a trigger and downstream targets drive the long-lasting defensive responses. Alternatively, prior exposure to shock, a necessary condition for rats to freeze upon silence onset, may lead to plastic changes that mediate a change from transient to sustained sound offset responses in any of the identified brain regions. Although we identified two input stations to the LA, there are multiple scenarios by which information may flow in this circuit. One scenario is that of a feed forward circuit whereby MGD drives VA, which in turn activates LA. This activation is probably indirect since evidence for direct projections from the VA to the amygdala is lacking^26^ (but see ^16^). Alternatively, feedback loops within the identified circuit may contribute to silence-triggered freezing. VA neurons project back to the MGD, allowing for top-down enhancement of relevant stimuli, possibly instructed by the amygdala. Indeed, neurons in LA that project back to primary auditory cortex are required for the expression of freezing to conditioned pure tones^27^. A similar feedback loop has been proposed for the encoding of behavioural relevance or context of visual stimuli, whereby the LA would contribute to the formation of neuronal ensembles in visual cortex that encode stimulus relevance, independent of ensembles that encode visual stimulus identity^28^.

The current study brings new insights into the mechanisms by which prey animals use the movement of others to infer danger, advancing our knowledge regarding how defence circuits respond to natural sound cues. By describing a pathway that includes auditory regions that have not been previously implicated in defence responses it expands the network that processes auditory cues in the context of danger. More broadly this study sets the stage to further our understanding of how sensory stimuli and their behavioural relevance are encoded by sensory systems and brain regions involved in motivated, survival behaviours.

## METHODS

### Animal Care

Naïve male Sprague Dawley rats (300–350 g) for *c-fos* experiments and 200-250g for optogenetic and muscimol experiments were obtained from a commercial supplier (Harlan and Charles Rivers, respectively). After arrival animals were pair housed in Plexiglas top filtered cages and maintained on a 12 h light/dark cycle with ad libitum access to food and water. All behavioural procedures were performed during the light phase of the cycle.

For the *c-fos* experiments, animals were kept in pairs and acclimated for at least one week before experimental manipulation. For optogenetic experiments animals were separated 4-6 days after arrival and subjected to virus injection and/or optic fibber implantation surgery. After this procedure animals were kept alone in Plexiglas boxes before experimental manipulation. For muscimol experiments animals were separated on the forth day after arrival and subjected to cannula implantation surgery. After the procedure, animals were kept alone for 7 days before experimental manipulations.

The Champalimaud Center for the Unknown follows the European Guidelines of animal care. The use of vertebrate animals in research in Portugal complies with the European Directive 86/609/EEC of the European Council.

### Viral vector, neuronal tracer and muscimol

The viral vector Adeno-associated virus expressing ArchT (AAV2/5 CAG-ArchT-GFP 1,3 ×10^12^ vg/ml) was produced by and purchased to University of North Carolina (UNC) vector core facility. The viral vector was diluted with sterile Phosphate Basal Solution (PBS) immediately before injection to obtain a concentration of 4 ×10^11^ vg/ml.

The transynaptic neuronal tracer Cholera Toxin subunit B Alexa Fluor 555 Conjugate (CT-B Alexa 555, 1 mg/mL) was produced by and purchased to Life Technologies. The GABA agonist muscimol (Cat# 2763-96-4) was bought to Sigma-Aldrich.

### Stereotactic surgery

Animals were anaesthetized with Isoflurane (Vetflurane 1000mg/g, Virbac) and placed in a stereotaxic apparatus (David Kopf Instruments). Small craniotomies were made using standard aseptic techniques.

For the Lateral Amygdala (LA) inhibition experiments, animals assigned to the ArchT+light and Control ArchT (Ct-ArchT) groups were targeted bilaterally to the LA (stereotaxic coordinates from Bregma, anterior–posterior: −3.3 mm, dorsal– ventral: −8.1, medial–lateral: 5.2 mm; derived from Paxinos and Watson 2007) using stainless steel guide cannula (24 gauge; Plastics One). Following cannula guide placement, 0.3 – 0.4 µL injections of rAAV2/5-CAG-ArchT-GFP were made through a stainless steel injection cannula (31 gauge; Plastics One), attached to a Hamilton syringe via polyethylene tubing. Injections were controlled through an automatic pump (PHD 2000; Harvard Apparatus). For animals assigned to the ArchT+light group optical fibbers (200µm, 0.37 numerical aperture, Doric lenses) were implanted in the LA (stereotaxic coordinates from Bregma, anterior–posterior: −3.3 mm, dorsal– ventral: −8.15 or 8.05 mm, medial–lateral: 5.2 mm) and affixed to the skull using stainless steel mounting screws (Plastics One, Inc.) and dental cement (TAB 2000, Kerr). No fibbers were implanted in animals assigned to the Ct-ArchT group. Rats were kept for 4 weeks before any behavioural manipulation to allow maximal expression of the virus. Animals of the Control light (Ct light) group were subjected to the same procedure, including implanting optical fibbers, but no virus was injected.

For the dorsal Medial Geniculate Nucleus (MGD) inhibition experiments, the procedures were similar but injections were targeted bilaterally to the MGD (stereotaxic coordinates from Bregma, anterior–posterior: −5.8 mm, dorsal– ventral: −5.3, medial–lateral: +/−3.4mm; derived from Paxinos and Watson 2007) and 0.2 µL of rAAV2/5-CAG-ArchT-GFP were injected.

For the ventral area (VA) and posteriodorsal (PD) inhibition experiments, animals were subjected to a stereotaxic surgery to bilaterally implant stainless steel guide cannulas (24 gauge, Plastics One) in either the PD (stereotaxic coordinates from Bregma, anterior-posterior −5.7 mm, dorsal-ventral −4.3mm and medial-lateral +6.5mm; derived from Paxinos and Watson 2007) or VA (stereotaxic coordinates from Bregma, anterior-posterior −3.5mm, dorsal-ventral −6.5mm, medial-lateral +6.5mm; derived from Paxinos and Watson 2007) areas of the Auditory Cortex. Following cannula implantation, a dummy cannula (31 gauge, Plastics One) was kept until test day.

For the retrograde tracing of auditory thalamus projections to this regions (used to confirm that the area that was inactivated indeed received input from the MGD) we took advantage of the already implanted guide cannulas, and posterior to the test day, CT-B Alexa 555 (0,2 µL) was bilaterally injected in half of the experimental animals following the same protocol as the one used to inject the viral vector. Animals were sacrificed 6 days after this procedure to allow sufficient transport of the tracer.

All injection sites and fibber placements were verified histologically and rats were excluded if either were not on the correct place.

### Behavioural Apparatus

Four distinct environments were used in this study - **Conditioning box**, **Test box - silence test**, **Tone conditioning box** and **Test box – tone test**.

Both **Conditioning boxes** (model H10-11R-TC, Coulbourn Instruments) have a shock floor of metal bars (model H10-11R-TC-SF, Coulbourn Instruments) and are placed inside a sound isolation chamber (Action automation and controls, Inc). In these boxes a precision programmable shocker (model H13-16, Coulbourn Instruments) delivered the foot-shocks, and tones were produced by a sound generator (RM1, Tucker-Davis Technologies) and delivered through a horn tweeter (model TL16H8OHM, VISATON).

The **Test box-silence test** consisted of a two-partition chamber made of clear Plexiglas walls (Gravoplot) divided in two equal halves, placed inside a sound attenuation chamber. The **Test box – tone test** consisted of a plastic box with a round platform in the bottom. Both **Test Boxes** had bedding on the floor. Tones were generated, delivered and calibrated using the same apparatus as the conditioning boxes. To minimize generalization between environments, conditioning with shock (Day 1) took place with light and Silence Test (Day 2) took place in the dark (Fig. 1a). For tone conditioning experiments, tone conditioning took place in the dark and recall took place under dim yellow light. Rats’ behaviour was tracked by a video camera mounted on the ceiling of the attenuating cubicle. A surveillance video acquisition system was used to record and store all videos on hard disk.

### Behavioural Procedures

All rats were pre-exposed to the different boxes prior to the beginning of the experiments. To test the role of auditory motion cues, the previously recorded movement-evoked sound (see^4^) was played during exposure to the Test box - silence test, and during the Silence Test.

#### Optogenetic experiments

For the **Silence test experiments**, rats were trained and tested as previously described^4^. During the test session, a fibber optic cable terminating in 2 ferrules (Branching Fibber-optic 200μm, 0.22 numerical aperture, Doric lenses) was connected to the chronically implanted optic fibbers. Animals in the ArchT+light and Ct light groups received laser illumination (estimated 30mW at the tip of the fibber) that started 10sec before the silence. Illumination lasted until 5 sec after the resumption of the playback of the movement-evoked sound. After the test session, animals returned to their home cages.

To test the role of the MGD in recall after tone conditioning, the same animals that were tested to the silence were exposed to the **Tone conditioning box** and **Test box – tone test** during the following four days of experiment. On the fifth day, after a 5-minute baseline animals received 3 tone-shock pairings (5Khz tone, 70dB, 15sec co terminating with1mA shock, 0.5sec). Animals were tested to the sound the day after in the Test box – tone test, where animals were exposed to 3 tone presentation, (5Khz tone, 70dB, 15sec). Laser illumination was identical to the one used in LA experiments.

#### *c-fos* experiments

Experiments followed a modified protocol of the Silence test experiments (described above), in which the test session lasted for 9 minutes. In these experiments, animals in the silence group were exposed to two periods of silence with the duration of one minute each (and with a period of 3 minutes separation) while for the control group (continuous sound) the recorded sound of movement was played throughout the entire test session.

#### Muscimol experiments

Experiments followed the same protocol as the **Silence Test Experiments**, however in the test day animals were injected with 0,2µl muscimol (0,5µg/µl concentration at 0,1µl/min rate) in either the VA or PD. The injections were performed with the help of an injection cannula, attached to a Hamilton syringe controlled by an automatic pump (PHD 2000; Harvard Apparatus). Animals were left undisturbed for 1h30min, period after which they were tested.

### Histology

Animals were deeply anesthetized with pentobarbital (600 mg/kg, i.p.) and transcardially perfused with PBS (0.01M), followed by ice-cold 4% paraformaldehyde (Paraformaldehyde Granular; cat#19210; Electron Microscopy Sciences) in 0.1 M phosphate buffer (PB) (PFA-PB). Brains were removed and postfixed in 4% PFA-PB and kept at 4ºC.

Coronal sections containing the LA (2.50mm to 3.80mm posterior to Bregma), VA (3.20mm to 4.60mm posterior to Bregma), PD (5.40mm to 6.7mm posterior to Bregma) and/or the MGB of the auditory thalamus (5.60mm to 6.50mm posterior to Bregma), were cut and mounted.

Mounted slices were rehydrated for 10 minutes with PBS and 500 µl of a 1:1000 diluted 4’, 6-diamidino-2-phenylindole (DAPI) solution (D9542, Sigma-Aldrich) was applied to each slide and incubated for 20 minutes in a shaker (10rpm). Slices were then washed 3 times with PBS and rinsed with distilled water. Slices were coversliped with Moviol 4-88 (81381, Sigma-Aldrich).

Fluorescent images were taken with the confocal microscope Zeiss LSM 710, Axioimager2, Objective Plan-Apochromat 20x/0.8M27 and software Zen 2010. Pictures were processed in ImageJ for compiling z-stacks, and exposure adjustments (same adjustment for both channels) were made with Photoshop.

Immunohistochemistry experiments examining *c-fos* expression in the MGB were performed in brain slices from animals that were rapidly and deeply anaesthetized with pentobarbital (600 mg/kg, i.p.) 2 h after the beginning of the behavioural paradigm. While anesthetized, animals were transcardially perfused with PBS, followed by ice-cold 4% PFA-PB. Brains were removed and postfixed in 4% PFA-PB for 24 h and subsequently cryoprotected in 20% glycerol (J.T.Baker) in 0.1 M PB for 72 h at 4ºC. Using a sliding microtome, sections of 40 µm containing the auditory thalamus (5.40 to 6.40 posterior to Bregma) were cut and collected in PBS. Next, sections were transferred to a 0.1% sodium azide (Sigma Aldrich) in PBS solution for storage. The immunohistochemical staining was performed simultaneously for all brain sections analysed. Staining was performed in free-floating sections. Sections were washed 3×10 min with PBS, incubated for 10 min with 0,9% H_2_O_2_ (Sigma Aldrich), washed again 3×10 min in PBS and blocked in PBS with1% bovine serum albumin (BSA) (cat#A7906, Sigma-Aldrich) and 0.1% Triton X-100 for 1h RT. Slices were then incubated O.N. RT with anti c-fos 1ry AB (1:500; rabbit; sc-52 Santa Cruz Biotechnology) in PBS with1% BSA and 0.1% Triton X-100. The next morning, sections were washed 3×10 min with PBS and incubated with goat anti-rabbit byotinilated 2ry antibody (1:1000; Cat#405008, Southern Biotec) in PBS with 1% BSA and 0.1% Triton X-100 for 1h RT. Sections were washed 3×10 min in PBS, incubated with Horseradish Peroxidase Streptavidin (1:300; cat#SA-5004; Vector Laboratories) in PBS with 0.2% Triton X-100 for 1h RT, washed 3×10 min in PBS-B and developed in diaminobenzidine tablets (DAB) (cat# D4418-50 SET; Sigma) for 3 min.

Sections were mounted on electrostatic slides, air dried, dehydrated in ethanol and xylenes and coverslipped with DPX. Brightfield images were taken in Zeiss Axioimager M1 microscope equipped with a CCD camera (Hamamatsu C8484), with objective 20x/0.80. Sections from comparable anteroposterior levels were selected for scoring and cell counts were scored using NIH Image J. For the initial analysis, cell counts for each subnucleus of the thalamus were averaged into a single score for each rat. For the analysis along the AP axis of the MGD we divided the sections in anterior (sections including and posterior to 5.64 until 5.76 (including) relative to Bregma), medial (sections posterior to 5.76 until 6.00 (including) posterior to Bregma) and posterior (sections posterior to 6.00 until 6.24 (including) relative to Bregma) and averaged the c-fos labelling cells in those slices.

### Analysis of Behavioural Data

Animals freezing behaviour during the Silence test was automatically scored using FreezeScan from Clever Sys. Baseline freezing levels were calculated using the median % of freezing during the 60s preceding the onset of silence. To avoid confounding factors such as freezing triggered by other cues that not the onset of silence, animals were considered outliers and thus excluded if their baseline freezing was higher than the 3^rd^ quart + 1.5 (3^rd^ quartile – 1^st^ quartile) baseline freezing of the population used in each experiment (LA, MGD, VA and PD experiments were considered separately). Due to the noise generated by the shutter used in the optogenetics experiments, animals that were freezing more than 50% in the 10sec period between the opening of the shutter and silence onset were also excluded.

### Statistical Analysis

For data analysis of the behavioural experiments we focused on the % of freezing during the silence gaps (that lasted one minute) and used as baseline the minute immediately preceding the silence interval. In this manner we ensure that both measures have the same sampling time.

A Shapiro-Wilk test was used to access the normality of our data. The behavioural data was in general not normally distributed and sample sizes were small, so we used non-parametric tests only.

For comparisons between groups (when comparing the change in freezing between baseline and silence periods) we used a Wilcoxon-Mann-Whitney test. For comparisons within group (comparing % of freezing during baseline and silence) we conducted a Wilcoxon signed-ranked test. For the *c-fos* behavioural data we averaged the percentage of freezing during the two periods of one minute preceding the silence inserts and the two periods of silence. For comparisons within the group (silence or continuous sound) we conducted a Wilcoxon signed-ranked test. For comparisons between groups of the *c-fos* labelled cells our data was not normally distributed and therefore we used a Mann-Whitney test. For analysing the effect of position along the AP axis and exposure to silence in the expression of *c-fos* we conducted a two-way ANOVA for unbalanced design given that our data was normally distributed. Statistical analysis was performed using MATLab.

## ACKNOWLEDGMENTS

We thank Alexandra Medeiros and Andreia Cruz for their input to the project, and Alexandra in particular for her effort in the histological characterization of the MGD. We also want to thank all the members of the Behavioral Neuroscience lab, as well as Samuel Walker and Tiago Marques, for comments on the manuscript.

This work was developed with the support from the Champalimaud Histopathology platform and the research infrastructure Congento LISBOA-01-0145-FEDER-022170. This project was supported by Fundação Champalimaud and ERC-2013-StG-337747

“C.o.C.O.”. Ana Pereira was supported by Fundação para a Ciência e Tecnologia, grant SFRH/BD/33943/2009.

## REFERENCES

1. Holle, L. I. & Radford, A. N. The development of alarm call behaviour in mammals and birds. Anim. Behav. 78, 791–800 (2009).

2. Murray, T. G., Zeil, J. & Magrath, R. D. Sounds of Modified Flight Feathers Reliably Signal Danger in a Pigeon. Curr. Biol. 27, 3520–3525 (2017).

3. Randall, J. A., Rogovin, K. A. & Shier, D. M. Antipredator behavior of a social desert rodent: footdrumming and alarm calling in the great gerbil, Rhombomys opiums. Behav Ecol Sociobiol 48, 110–118 (2000).

4. Pereira, A. G., Cruz, A., Lima, S. Q. & Moita, M. A. Silence resulting from the cessation of movement signals danger. Curr. Biol. 22, 627–628 (2012).

5. Catania, K. C., Hare, J. F. & Campbell, K. L. Water shrews detect movement, shape, and smell to find prey underwater. Proc. Natl. Acad. Sci. 105, 571–576 (2008).

6. Carr, C. E. & Christensen-Dalsgaard, J. Sound localization strategies in three predators. Brain. Behav. Evol. 86, 17–27 (2015).

7. Friedel, P., Young, B. A. & Van Hemmen, J. L. Auditory localization of ground-borne vibrations in snakes. Phys. Rev. Lett. 100, 2–5 (2008).

8. Zhao, Z. et al. Zona incerta GABAergic neurons integrate prey-related sensory signals and induce an appetitive drive to promote hunting. Nat. Neurosci. 22, 921–932 (2019).

9. Mortimer, B., Rees, W. L., Koelemeijer, P. & Nissen-meyer, T. Classifying elephant behaviour through seismic vibrations. Curr. Biol. 28, R547–R548 (2018).

10. Herry, C. & Johansen, J. P. Encoding of fear learning and memory in distributed neuronal circuits. Nat. Neurosci. 17, 1644–1654 (2014).

11. Theunissen, F. E. & Elie, J. E. Neural processing of natural sounds. Nat. Rev. 15, (2014).

12. Moczulska, K. E. et al. Dynamics of dendritic spines in the mouse auditory cortex during memory formation and memory recall. Proc. Natl. Acad. Sci. 110, 18315–18320 (2013).

13. Atsak, P. et al. Experience modulates vicarious freezing in rats: A model for empathy. PLoS One 6, (2011).

14. Wiegert, J. S., Mahn, M., Prigge, M., Printz, Y. & Yizhar, O. Silencing Neurons: Tools, Applications, and Experimental Constraints. Neuron 95, 504–529 (2017).

15. Doron, N. N. & Ledoux, J. E. Cells in the posterior thalamus project to both amygdala and temporal cortex: A quantitative retrograde double-labeling study in the rat. J. Comp. Neurol. 425, 257–274 (2000).

16. Romanski, L. & LeDoux, J. Information Cascade from Primary Auditory Cortex to the Amygdala: Corticocortical and Corticoamygdaloid Projections of the Temporal Cortex in the Rat. Cereb. Cortex 515–532 (2006).

17. Bordi, F. & LeDoux, J. E. Response properties of single units in areas of rat auditory thalamus that project to the amygdala. Exp. brain Res. 98, 261–274 (1994).

18. He, J. ON and OFF pathways Segregated at the Auditory Thalamus of the Guinea Pig. 21, 8672–8679 (2001).

19. Smith, P. H., Uhlrich, D. J., Manning, K. A. & Banks, M. I. Thalamocortical projections to rat auditory cortex from the ventral and dorsal divisions of the medial geniculate nucleus. J. Comp. Neurol. 520, 34–51 (2012).

20. Orsini, C. A. & Maren, S. Glutamate receptors in the medial geniculate nucleus are necessary for expression and extinction of conditioned fear in rats. Neurobiol. Learn. Mem. 92, 581–589 (2009).

21. Donishi, T., Kimura, A., Okamoto, K. & Tamai, Y. ‘Ventral’ area in the rat auditory cortex: A major auditory field connected with the dorsal division of the medial geniculate body. Neuroscience 141, 1553–1567 (2006).

22. Liu, J. et al. Parallel Processing of Sound Dynamics across Mouse Auditory Cortex via Spatially Patterned Thalamic Inputs and Distinct Areal Intracortical Circuits. Cell Rep. 27, 872–885.e7 (2019).

23. Llano, D. A. & Sherman, S. M. Evidence for Nonreciprocal Organization of the Mouse Auditory Thalamocortical-Corticothalamic Projection Systems. J. Comp. Neurol. 507, 1209–1227 (2008).

24. Doron, N. N., Ledoux, J. E. & Semple, M. N. Redefining the tonotopic core of rat auditory cortex: Physiological evidence for a posterior field. J. Comp. Neurol. 453, 345–360 (2002).

25. Kopp-Scheinpflug, C., Sinclair, J. L. & Linden, J. F. When Sound Stops: Offset Responses in the Auditory System. Trends Neurosci. 41, 712–728 (2018).

26. Kimura, A., Donishi, T., Okamoto, K., Imbe, H. & Tamai, Y. Efferent connections of the ventral auditory area in the rat cortex: Implications for auditory processing related to emotion. Eur. J. Neurosci. 25, 2819–2834 (2007).

27. Yang, Y. et al. Selective synaptic remodeling of amygdalocortical connections associated with fear memory. Nat. Neurosci. 19, 1348–1355 (2016).

28. Ramesh, R. N., Burgess, C. R., Sugden, A. U., Gyetvan, M. & Andermann, M. L. Intermingled Ensembles in Visual Association Cortex Encode Stimulus Identity or Predicted Outcome. Neuron 100, 900–915.e9 (2018).

