## Supplementary material for "A new thalamo-cortical-amygdala circuit is involved in processing a natural auditory alarm cue": Suplementary Figures

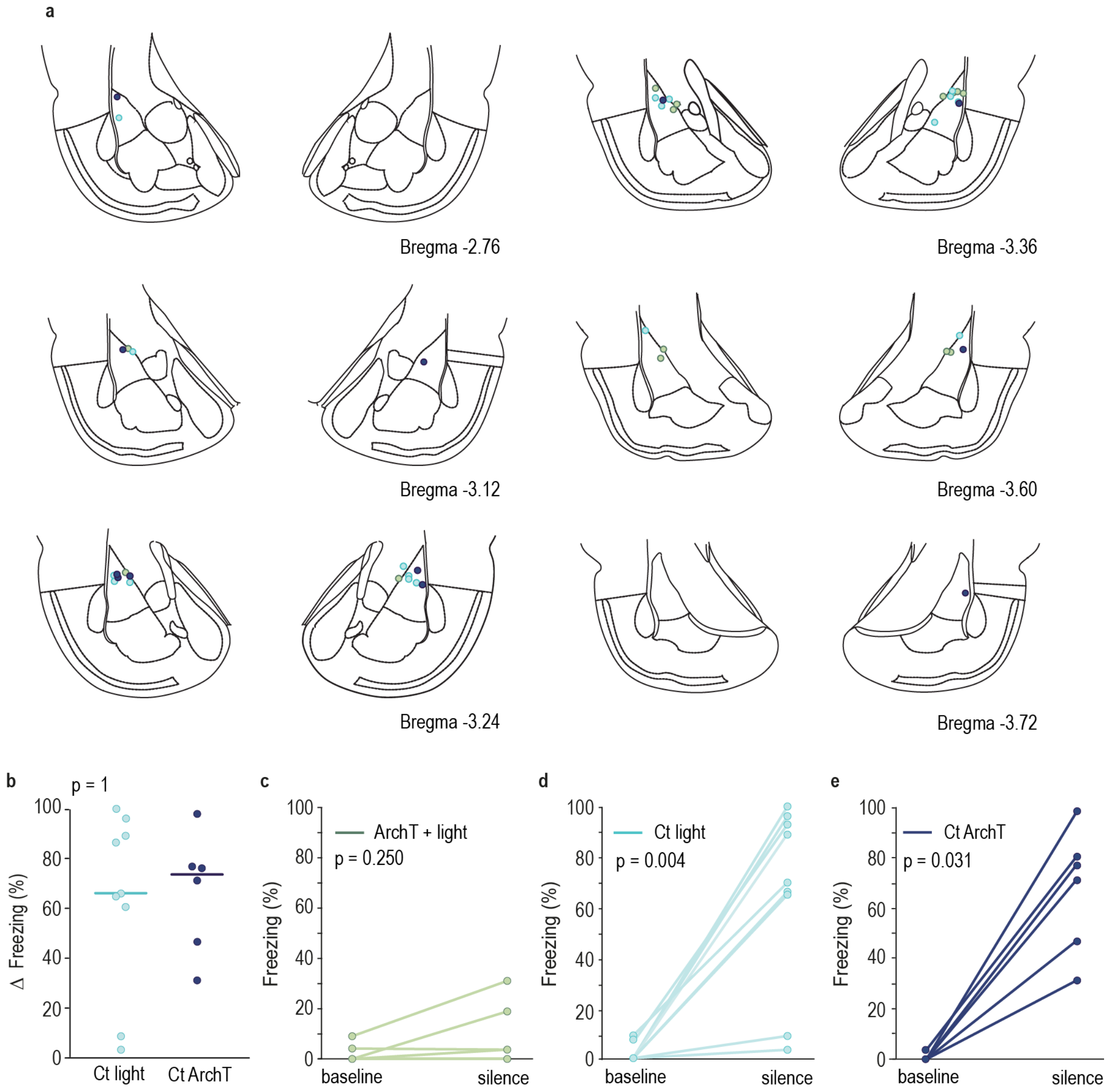


**Supplementary Figure 1. The LA is necessary for the expression of freezing driven by the cessation of movement-evoked sound. a,** Coronal slices representing fiber placements and/or injection site for the ArchT + light (light green), Control Light (light blue) and Control ArchT (dark blue) groups. **b,** Same as Fig.1 d, but for animals of the 2 control groups. Horizontal bar represents the median value of the group (Control light = 66.13%, Control Virus = 73.67%, Wilcoxon Rank Sum). **c, d,** and **e,** Line graph showing average time spent freezing during the minute immediately preceding the cessation of the movement-evoked sound (baseline) and the minute of silence for each rat of the ArchT + light (n=7), Ct light (n=9) and Ct ArchT (n=6) groups. Wilcoxon Signed Rank Test.

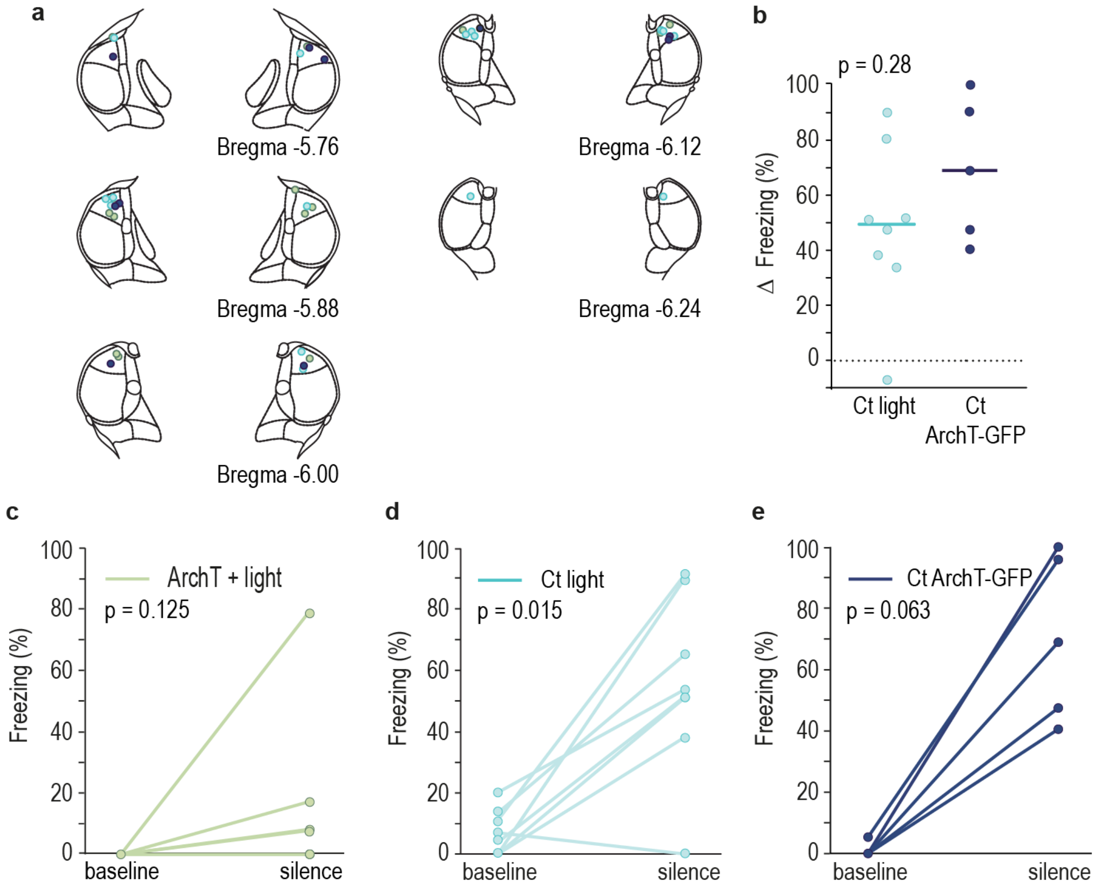


**Supplementary Figure 2. The MGD is necessary for the expression of freezing driven by the cessation of the movement-evoked sound but not by a discrete auditory cue. a,** Coronal slices representing fiber placements and/or injection site for the ArchT + light (light green), Control light (light blue) and Control ArchT (dark blue) groups. **b,** Same as Supplementary Fig. 1b but for rats with surgeries targeting the MGD. Horizontal bar represents the median value (Ct light = 49.47%, Ct ArchT = 68.93%, Wilcoxon Rank Sum test, ranksum = 48). **c, d,** and **e,** Same as Supplementary Figure 1 c, d, and e, but for rats of the ArchT + light (n=7), Control light (n=8) and Control ArchT (n=5) groups with surgeries targeting the MGD. Wilcoxon Signed Rank Test, baseline *vs* silence ArchT + light signedrank = 0; Control light signedrank = 1; Control ArchT signedrank = 0

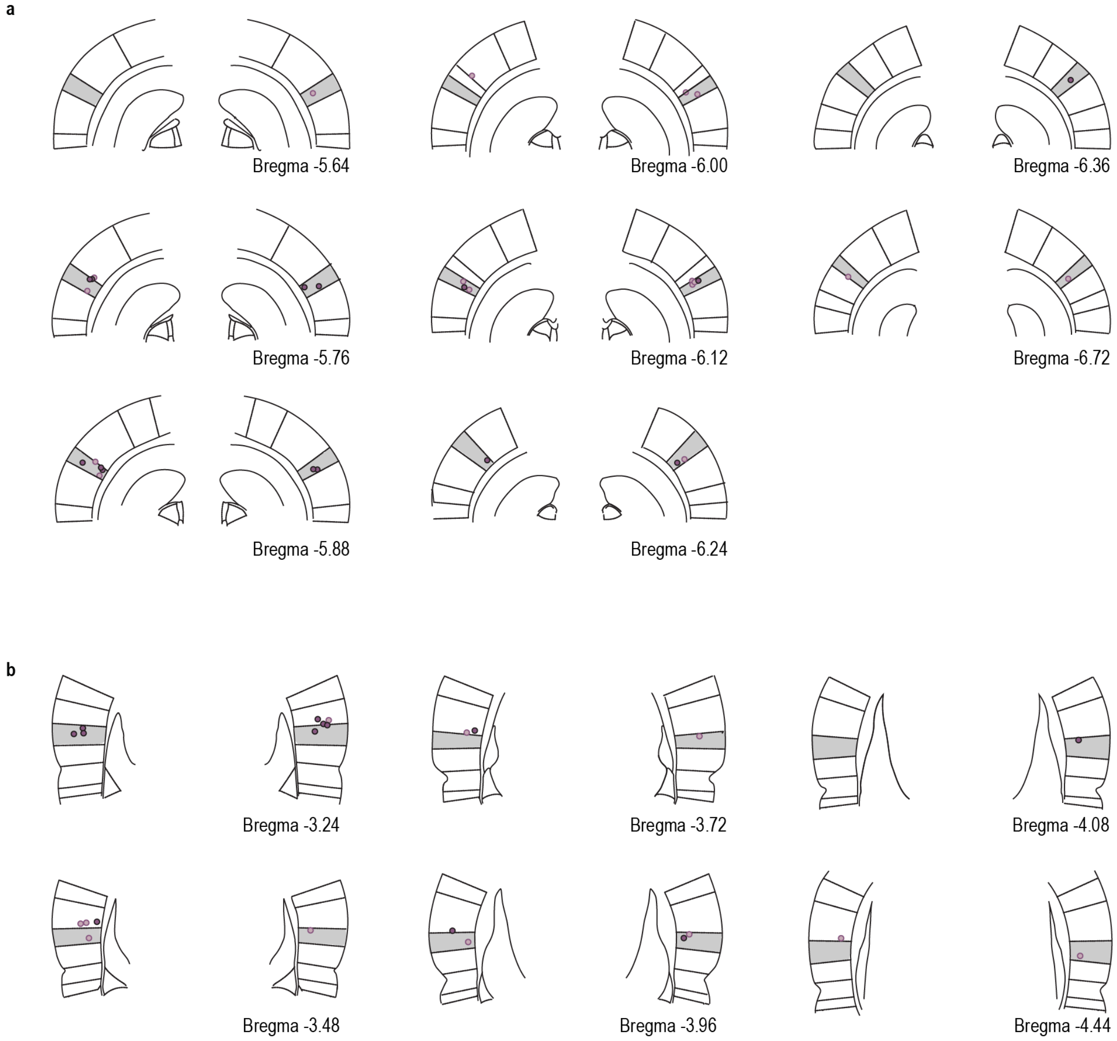


**Supplementary Figure 3. Coronal slices representing injection site** for PBS (light purple) and Muscimol (dark purple) in **a,** posterodorsal (PD) and **b,** ventral area (VA). Areas shaded in grey correspond to PD and VA (respectively).
